# LAMA5 deficiency disrupts ECM–WNT crosstalk in chondrogenesis and contributes to idiopathic short stature

**DOI:** 10.1101/2025.11.19.689218

**Authors:** Alexander Schulz, Emily M. Brockmann, Steffen Uebe, Arif B. Ekici, Christian T. Thiel

## Abstract

Idiopathic short stature (ISS) affects 2%–3% of the population and is genetically heterogeneous, with emerging evidence implicating the extracellular matrix (ECM) of the growth plate. We identify *LAMA5*, encoding laminin-α5, as a candidate ISS gene, with rare heterozygous variants present in 1.2% of affected individuals. To define its functional role, we generated CRISPR/Cas9-mediated LAMA5-knockout (KO) urine-derived stem cells (USCs) and induced chondrogenic differentiation in two- and three-dimensional culture systems. Loss of LAMA5 impaired chondrogenesis, with disruption of cell–cell junction programs and abnormal architecture of chondrogenic spheroids. Bulk RNA sequencing combined with weighted gene co-expression network analysis revealed *WNT7A* and *FLI1* as key dysregulated genes within the module most strongly associated with the KO phenotype. Gene Ontology enrichment of this module highlighted embryonic limb morphogenesis as the top biological process, and WNT7A was assigned to canonical WNT signaling. Pharmacologic activation of WNT signaling using lithium chloride (LiCl) partially restored expression of *WNT7A*, *FLI1*, *TFAP2A*, *GRHL2*, and *PITX1* toward wild-type levels, indicating that attenuated WNT activity is a principal downstream consequence of LAMA5 deficiency. Consistent with this, we identified an individual with ISS carrying a heterozygous *PITX1* missense variant, supporting convergence of ECM (LAMA5) and transcriptional (PITX1) perturbations on a shared WNT-centered limb-morphogenesis network. Together, these findings demonstrate that laminin-α5 is required for proper ECM–WNT signaling integration during human chondrogenesis and suggest that dysregulated WNT activity represents a mechanistic link between LAMA5 dysfunction and impaired endochondral growth. Partial rescue by WNT pathway re-activation highlights a potentially targetable downstream mechanism in ISS pathogenesis.

## Introduction

Idiopathic short stature (ISS) is a common pediatric growth disorder, affecting approximately 2–3% of children, and is defined as a height more than two standard deviations below the population mean in the absence of identifiable endocrine, nutritional, or systemic causes^1,2^. Its genetic architecture is highly heterogeneous, ranging from polygenic contributions with minor effect size on growth to rare, high-impact variants underlying monogenic severe cases^3–7^. Recent genome sequencing studies have increasingly implicated the extracellular matrix (ECM) of the growth plate as a critical mediator of linear growth, with genes such as *ACAN*, *COL2A1*, and *MATN3* underscoring the importance of the chondrocyte microenvironment in organizing proliferation, hypertrophy, and cartilage organization^8–12^.

Laminins, a major part of the ECM, are a family of heterotrimeric glycoproteins that constitute major structural and functional components of basement membranes, providing not only mechanical support but also instructive cues for tissue morphogenesis^13^. They regulate cell adhesion, polarity, survival, and differentiation through interactions with integrins, dystroglycans, and other ECM receptors, integrating extracellular architecture with intracellular signaling networks^14–18^. Among the laminin isoforms, laminin α5, encoded by *LAMA5*, is broadly expressed during organogenesis and enriched in the pericellular matrix of developing cartilage^19–21^. In humans, biallelic mutations in□*LAMA5*□cause a severe skeletal dysplasia (MIM#620076) characterized by bent long bones, delayed ossification and disrupted focal⍰adhesion kinase signaling in chondrocytes, highlighting a non⍰redundant role for laminin□α5 in skeletal morphogenesis^19^.□

Despite this evidence, the precise mechanisms by which laminin α5 modulates human chondrogenesis, and its potential interaction with canonical growth-regulatory pathways such as WNT and BMP signaling^22–25^, remain poorly defined. Understanding these mechanisms is critical to elucidate how ECM perturbations contribute to ISS, particularly as the growth plate pericellular microenvironment functions as both a structural scaffold and a signaling hub that coordinates chondrocyte proliferation, hypertrophy, and matrix deposition^12,26–29^.

Based on the identification of rare heterozygous *LAMA5* variants in 1.2 % of our cohort of individuals with ISS^30^, this study employs a patient-relevant human stem cell model of chondrogenesis to dissect the functional consequences of LAMA5 deficiency. Using a multi-scale approach that integrates three-dimensional spheroid morphogenesis, transcriptomic profiling, and gene network analyses, we aim to determine how LAMA5 loss alters cellular architecture, adhesion programs, and developmental signaling pathways. The overarching goal is to define the mechanistic link between laminin α5 function, ECM integrity, and transcriptional networks governing endochondral bone growth, providing insight into the molecular basis of ISS and identifying potential targets for therapeutic intervention.

## Methods

### Urine stem cell cultivation and generation of LAMA5-KO USC

USCs were isolated and expanded as described previously^31^. To summarize, urine was centrifuged (300 g) and cell pellets washed. USCs were cultured and expanded after repeated centrifugation (300 g) in culture Media (1:1 mixture of DMEM high glucose (Gibco) and KSFM (Gibco) with additives).

To induce frameshift mutations, we used the Alt-R™ CRISPR-Cas9 System (Integrated DNA Technologies) according to the manufacturer, including predesigned Alt-R CRISPR crRNAs, tracrRNAs (ATTO 550™), Lipofectamine CRISPRMAX (Invitrogen) and TrueCut Cas9-Protein v2 (Invitrogen). Components were mixed in Optimem (Gibco), the recommended amount of transfected ribonucleoprotein complex was doubled and antibiotics removed during transfection period (approximately 18 hours). Transfected cells (ATTO550^+^), were FACS sorted the next day in a density of one cell per well, utilizing rhLaminin-521 (Gibco) coating for expansion and application of Y-27632 (BD Biosciences) for 24 hours (Figure S1A). The Laminin coating was removed prior to comparative experiments.

### Validation of LAMA5-KO clones

To validate genomic cut at the gRNA target site of individual clones, we amplified the region of interest with Platinum™ Direct PCR Universal Master Mix (Invitrogen), from the supernatant of cooked cells (95 °C, 10 minutes). This was followed by standard clean up protocols and sanger sequencing with BigDye Terminator Kit v3.1 (Applied Biosystems). Clones with homozygous deletions or insertions were expanded. Three established KO cell lines with different frameshift inducing deletions (KO1, KO2, KO3) were subjected to RT-qPCR, western blot analysis and bulk RNA-Seq.

### Differentiation of USCs and alcian blue / alizarin red staining

USCs were differentiated (osteogenic, chondrogenic) as already outlined^31^. In brief, USCs were cultured in osteogenic^32^ or chondrogenic (StemPro, Gibco) media for two weeks and then stained for GAGs (alcian blue) or calcium deposits (alizarin red).

### Bulk RNA-Sequencing and differential gene expression analysis

We processed Bulk RNA-seq data as detailed in our previous publication^31^. USC samples were analyzed using DESeq2 (v1.48.1)^33^ for differential expression (between cell types: osteogenic, chondrogenic, undifferentiated; between genotypes: wildtype and knockout), with significance defined at adjusted p < 0.05. Gene annotations were mapped to HGNC symbols, and results were visualized. Variance-stabilized data (rlog) were filtered for low variance, and PCA was performed using prcomp(). For volcano plot visualisation we used the R package ggplot2 (v3.5.2)^34^, for heatmaps pheatmap (v1.0.13)^35^, supported by ggrepel (v0.9.6)^36^ and RColorBrewer (v1.1.3)^37^.

### Functional analysis of LAMA5-KOs

RNA was extracted with RNeasy Mini Kit (Qiagen) with DNAse digestion or NucleoSpin RNA Plus (Machery Nagel) with gRNA removal column, according to manufacturers. For cDNA synthesis we used the LunaScript RT SuperMix Kit (New Englang Biolabs). RT-qPCR of cDNA was performed using standard TaqMan assay protocols (Applied Biosystems). We used four technical, and at least three biological replicates per analysis. Analysis (RQ calculation, unpaired welch’s t-test of ddCT values, visualization) was performed in R (v4.5.1) and Graphpad Prism (v10).

For western blotting we used 4–15% Mini-PROTEAN TGX (Bio-Rad Laboratories) for separation of proteins (300 V, 30 minutes). We transferred proteins onto nitrocellulose membrane with a semi-dry blot chamber (Trans-Blot Turbo Transfer System, Bio-Rad Laboratories, 1.3 A, 18 minutes) in towbin transfer buffer. Then we used the SuperSignal™ Western-Blot-enhancer (Thermo Scientific), according to the manufacturer’s manual. Mouse anti-LAMA5 antibody (Atlas Antibodies, AMAb91124) was used in a dilution of 1:1000, secondary goat anti-mouse antibody-HRP (Invitrogen, 31430), in a dilution of 1:10000 in 5 % milk powder (Roth) in TBST buffer. Sequentially, rabbit anti-Beta actin (abcam, ab8227), was incubated overnight at 4 °C in a dilution of 1:1000 in TBST, secondary goat anti-rabbit-HRP (Cell Signaling, 7074) was used in a concentration of 1:10000 in TBST. Detection was performed with the SuperSignal™ West Femto Maximum Sensitivity substrate (Thermo Scientific). (Raw images: Figure S3B-C).

For spheroid formation, we seeded 20000 USCs in each well of a 96 well Nunclon Sphera ultra low attachment cell culture plate (ThermoFisher Scientific), followed by centrifugation for 10 minutes at 300 g. Culture lasted for 3 weeks with StemPro Chondrogenesis Kit (Gibco). Fixation was done for one hour with 4 % paraformaldehyde (PFA) at room temperature, followed by permeabilization with Triton X-100 for 30 min and blocking with 1 % bovine serum albumin (Sigma-Aldrich) in PBS for 45 minutes. For primary antibody incubation anti-aggrecan (abcam, ab3778) and anti-LAMA5 (abcam, ab220837) were used in a concentration of 1:500 for one hour. Secondary anti-mouse antibody (AlexaFluor 488, Invitrogen) and anti-rabbit antibody (AlexaFluor 594, Invitrogen) were incubated for one hour in a 1:500 dilution, with addition of DAPI (1:1000, thermo fisher scientific). Spheroids were mounted with aqua polymount (Polysciences). Pictures (Z-stack projections of 25 pictures) were taken with the same exposure times (ACAN: 35 ms, LAMA5: 30 ms, DAPI 10 ms). The same image enhancements for each condition and size/circularity measurements of spheroids were performed with FIJI^38^. The macro used for size measurements will be provided in our Github repository upon publication.

### Reactome pathway enrichment and STRING network analysis

Differentially expressed genes (DEGs) were filtered at adjusted p-value (padj) < 0.05 and |log2 fold-change| > 1. ENSEMBL identifiers were mapped to Entrez gene IDs using biomaRt (v2.64.0)^39^. Reactome pathway enrichment was carried out with enrichPathway() (organism = human, p < 0.05) in ReactomePA (v1.52.0)^40^. Enrichment results were visualized using enrichplot (v1.28.4)^41^, ggplot2 (v3.5.2)^34^, and RColorBrewer (v1.1.3)^37^. A category–gene network (cnetplot) was constructed for Reactome pathways.

For network-level analysis, genes from the Reactome pathway Cell junction organization (R-HSA-446728) were retrieved via reactome.db (v1.92.0)^42^ and mapped to DEGs. High-confidence protein–protein interactions (confidence score ≥ 0.7) among these genes were obtained from STRING (v11) through the STRINGdb (v2.20.0)^43^ package. An undirected network was constructed in igraph (v2.1.4)^44^, annotated with log2 fold-change values, and pruned by excluding nodes with |log2FC| < 1 and retaining only the largest connected component. Node sizes were scaled to |log2FC|, node colors reflected expression changes, and edge widths represented STRING combined confidence scores. The final network was visualized using ggraph (v2.2.1)^45^ with a force-directed layout and labels added via ggrepel (v0.9.6)^36^.

### Weighted gene coexpression network analysis

Weighted gene co-expression network analysis was performed in R using WGCNA (v1.73.0)^46^. Raw counts were transposed to samples × genes, filtered to retain genes with counts >1 in ≥4 samples, and log₂(count+1)-transformed. Soft-thresholding (powers 1–20) was evaluated with pickSoftThreshold() and a power of 9 was chosen (Figure S1C). Networks were constructed with blockwiseModules() (power = 9, TOMType = “unsigned”, minModuleSize = 30, reassignThreshold = 0, mergeCutHeight = 0.25, numericLabels = TRUE, pamRespectsDendro = FALSE) with multithreading enabled (allowWGCNAThreads()). Module eigengenes were computed (moduleEigengenes()), correlated with sample traits (KO vs WT and cell type) using Pearson correlation and corPvalueStudent() for p-values, and module membership (kME) was calculated as gene–eigengene Pearson correlations; hub genes were defined as the top 15 by |kME|. For modules of interest (e.g., “lightgreen”), Ensembl to Entrez/HGNC mapping was performed with biomaRt (v2.64.0)^39^ and GO enrichment tested with clusterProfiler (v4.16.0)^47^ using BH correction (q < 0.05). Visualizations employed pheatmap (v1.0.13)^35^, ComplexHeatmap (v2.24.1)^48^ and circlize (v0.4.16)^49^ for heatmaps, ggplot2 (v3.5.2)^34^ and enrichplot (v1.28.4)^41^ for enrichment plots, and igraph (v2.1.4)^44^ or ggraph (v2.2.1)^45^ with ggrepel (v0.9.6)^36^ for module network visualizations; module TOMs were thresholded at 0.25 to derive weighted igraph objects and key genes (e.g., FLI1, WNT7A) were highlighted.

### Lithium chloride treatment of LAMA5-KO cells

Replicates of the LAMA5-KO2 cell line were seeded directly or on LM521-coated plates (10 µg/ml) in parallel to WT USCs and cultured for 7 days. For WNT pathway activation, some replicates (n = 4) were treated with 20 mM LiCl during the final 24 hours. FLI1 expression was assessed by RT-qPCR. For global transcriptional analysis, KO2 ± LiCl samples were subjected to bulk RNA-sequencing. KO2 + LiCl samples were combined with previous WT and untreated KO2 counts, variance-stabilized using DESeq2, and batch-corrected with limma::removeBatchEffect. A batch-corrected rescue index was calculated for each gene as (KO2+LiCl – KO2)/(WT – KO2), where 1 indicates full restoration to WT expression, and visualized as bar plots for hub genes and key limb morphogenesis genes. To complement these analyses, raw gene expression (TPM) was also computed without batch correction, and WT vs KO or KO vs KO+LiCl comparisons were plotted with error bars, individual sample points, and significance annotations. Gene identifiers were mapped to HGNC symbols or manually curated aliases where necessary, and plots were gunenerated with ggplot2 (v3.5.2) and patchwork (v1.3.2)^50^.

### PITX1 structural modelling and visualization

Wild-type (WT) and mutant (p.M205L) PITX1 structural models were generated using ColabFold/AlphaFold2 (Sokrypton ColabFold notebook: https://colab.research.google.com/github/sokrypton/ColabFold/blob/main/AlphaFold2.ipynb) with default parameters. Input sequences were identical except for a single amino acid substitution introduced in the FASTA prior to prediction. Resulting model files (top ranked models) were downloaded and inspected in PyMOL v3.1.6.1 (Schrödinger, LLC). Structures were rigid-superposed using PyMOL’s super command. For visualization and domain annotation we colored the canonical DNA-binding homeodomain (residues 89–148) green, the OAR motif (residues 280–293) red, and the mutated residue 205 cyan. Distances between equivalent Cα atoms (residue 205) were measured with the distance command and dashed distance objects were annotated in the figures.

## Results

### Generation and validation of LAMA5-knockout USC lines

To directly assess the contribution of laminin α5 (LAMA5) to human skeletal differentiation, we established LAMA5-deficient urine-derived stem cell (USC) lines using CRISPR/Cas9-mediated gene editing. Three independent LAMA5-knockout (KO) clones were generated, each harboring homozygous disruptive frameshift alleles confirmed by Sanger sequencing. Additionally, reduced expression was validated with RT-qPCR and absence of protein verified with western blot analysis (Figure 1A). This ensured that any downstream phenotypes could be confidently attributed to the absence of laminin α5, rather than incomplete editing or clonal variability.

**Figure 1:**
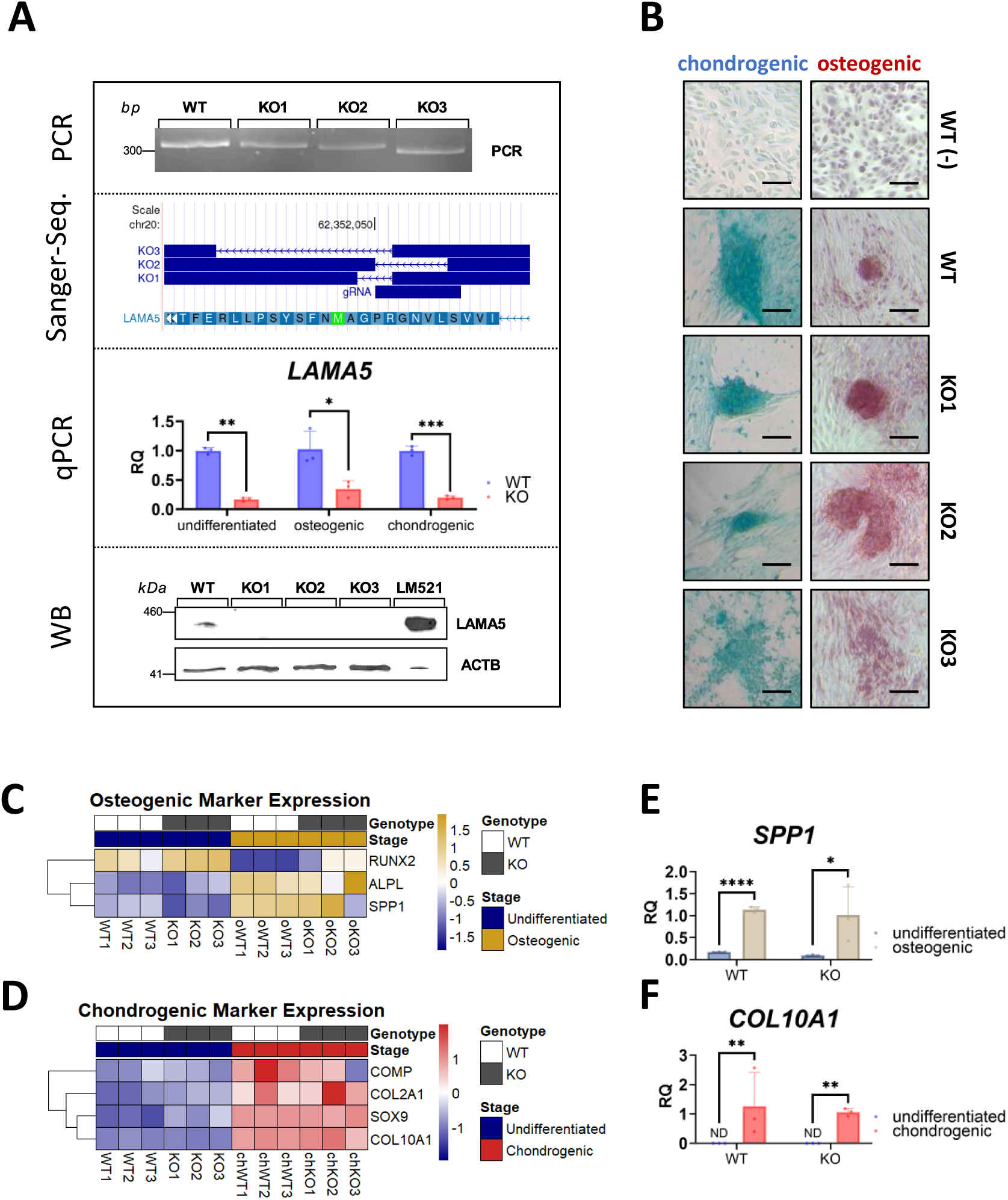
Generation and differentiation capacity of LAMA5-KO urine-derived stem cells (USCs). (A) PCR, sanger sequencing, RT-qPCR and western blot confirming frameshift-inducing indels in three independent clones (KO1–KO3). Unedited western blots are in Figure S3. (B) Representative alcian blue (glycosaminoglycans) and alizarin red (mineralization) staining of WT (from previous publication^31^) and LAMA5-KO USCs after chondrogenic and osteogenic differentiation (scale bars = 100 µm). (C) Expression of canonical osteogenesis associated genes as determined by bulk RNA-Seq. (D) Expression of canonical chondrogenesis associated genes as determined by bulk RNA-Seq. (E) Quantification by RT-qPCR of selected late osteogenic differentiation associated gene: *SPP1*, (RQ undifferentiated WT = 0.17 ± 0.01, RQ differentiated WT = 1.14 ± 0.06, p = 2 x 10^-6^, Welch’s t-test, n = 3; RQ undifferentiated KO = 0.09 ± 0.01, RQ differentiated KO = 1.02 ± 0.64, p = 0.0264, Welch’s t-test, n = 3). (F) Quantification by RT-qPCR of selected late chondrogenic differentiation associated gene: *COL10A1*, (RQ undifferentiated WT = 0.0006 ± 0.0003, RQ differentiated WT = 1.2675 ± 1.1514, p = 0.0019, Welch’s t-test, n = 3; RQ undifferentiated KO = 0.0012 ± 0.0006, RQ differentiated KO = 1.0605 ± 0.1298, p = 0.0018, Welch’s t-test, n = 3).

Because laminin α5 is a key extracellular matrix (ECM) component, its loss could nonspecifically impair cell viability or multipotency. To exclude these confounders, we first assessed the baseline differentiation potential of wild-type (WT) and LAMA5-knockout (KO) USCs. Both genotypes retained the ability to undergo osteogenic and chondrogenic differentiation, as evidenced by Alizarin Red and Alcian Blue staining for calcium and glycosaminoglycan deposition, respectively (Figure 1B). This confirmed that LAMA5 loss does not in general compromise lineage commitment or ECM secretion.

Transcriptomic profiling was then performed to further substantiate these findings. Bulk RNA-sequencing revealed upregulation of differentiation associated genes in both WT and LAMA5-KO cells (Figure 1 C, D), indicating successful lineage specification at the global level. To validate this and to assess terminal differentiation, RT–qPCR was performed for mature chondrogenic and osteogenic gene expression. Expression of *COL10A1* and *SPP1*, genes associated with terminal chondrocytes and osteoblasts, were robustly induced upon lineage induction in both genotypes (Figure 1E, F). These results confirm that LAMA5-KO USCs not only enter chondrogenic and osteogenic lineages but can also activate late-stage differentiation programs. Principal component analysis (PCA) showed clear segregation by both genotype and differentiation state (Figure S1B), underscoring the reproducibility and biological distinctness of the model.

Together, these results establish a valid experimental platform: LAMA5-KO USCs remain viable and lineage-competent, allowing mechanistic dissection of laminin α5’s specific role in morphogenesis and signaling without confounding cytotoxic effects.

### LAMA5 deficiency induces a junctional transcriptional signature during chondrogenesis

After confirming the differentiation potential of LAMA5-deficient USCs, we next sought to identify molecular programs disrupted by its loss during chondrogenesis. Differential gene expression analysis between WT and KO chondrogenic cultures revealed striking downregulation of *CDH1* (E-cadherin) in KO cells (Figure 2A). As cadherins mediate adherens junctions critical for tissue cohesion, its reduction suggested altered intercellular communication and polarity.

**Figure 2:**
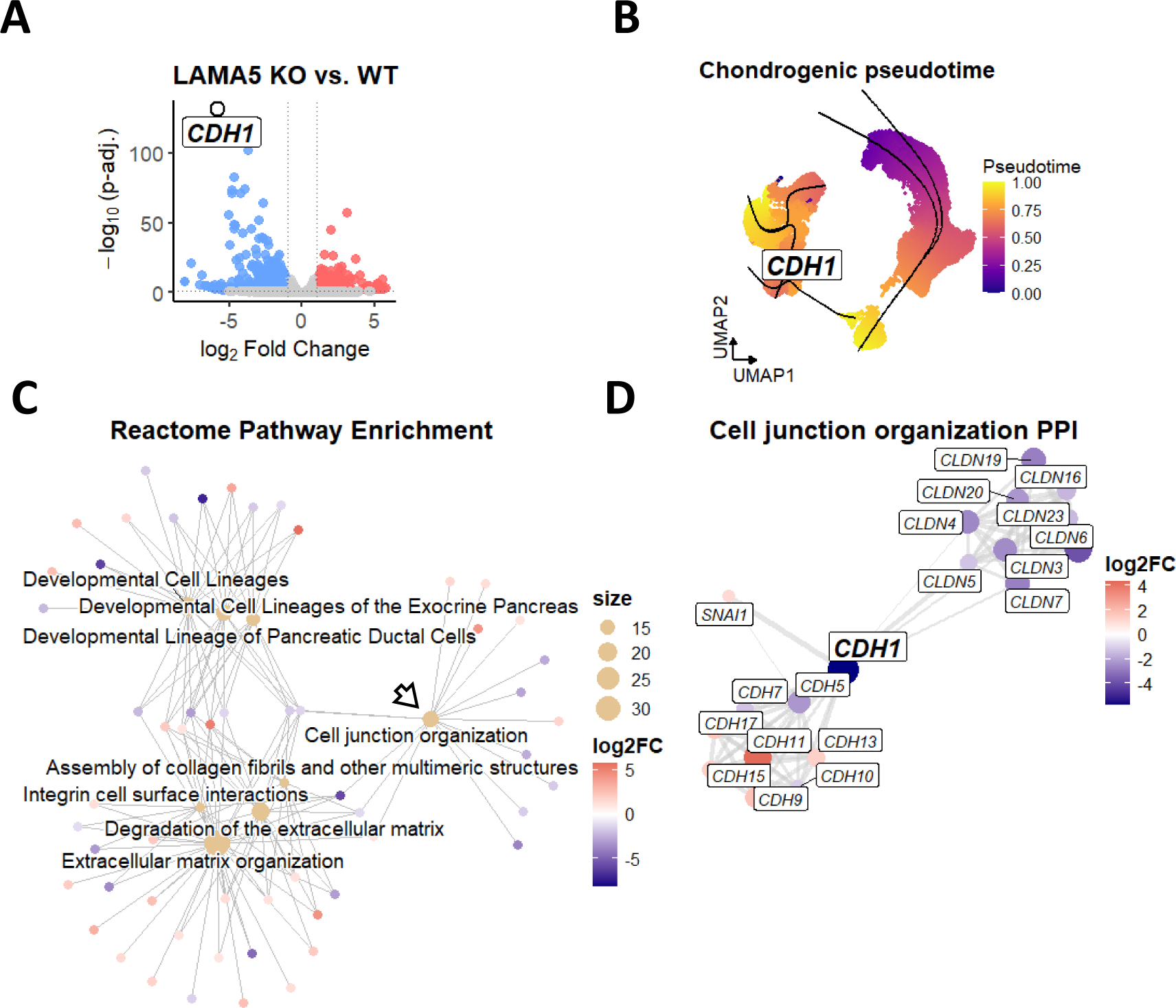
LAMA5 loss impairs chondrogenesis via disruption of cell junction organization. (A) Volcano plot showing strong downregulation of *CDH1* in KO chondrogenic cells (log2FC = −5.79; adjusted p = 5.09×10⁻¹³²; n = 3). (C) (B) *CDH1* expression across chondrogenic differentiation in single cell RNA seq pseudo time (data modified from Schulz et al.^31^). (C) Reactome pathway enrichment for DEGs in chondrogenic KO versus WT. The arrows highlights “Cell junction organization”, a key deregulated pathway which genes were investigated in downstream analysis (displayed categories = top 8). (D) STRING protein–protein interaction network for genes annotated to the Cell junction organization pathway; nodes are scaled by |log2FC| and colored by direction of change (blue = down, red = up).

To interpret this within a developmental framework, we mapped *CDH1* expression onto a published single-cell pseudotime atlas of chondrogenesis^31^. In normal differentiation, *CDH1* expression peaks at late pseudotime, corresponding to mature chondrogenic stages (Figure 2B). The observed reduction in KO cells therefore reflects a failure to activate a late-stage adhesion program in chondrogenesis.

Pathway enrichment analysis of differentially expressed genes was done to reinforce this interpretation, highlighting “Cell junction organization” among the top enriched Gene Ontology (GO) terms (Figure 2C). We then constructed a STRING network from the genes of this GO-term, to identify impaired functional connections within this pathway. *CDH1* at the network core connected with other differentially expressed cadherins, with coordinated downregulation of several *CLDN* (claudin) family members (Figure 2D). Since claudins are structural components of tight junctions, these data collectively indicate that *LAMA5* deficiency selectively disturbs the transcriptional program responsible for both adherens and tight junction assembly.

This therefore establishes a key mechanistic link, highlighting that laminin α5 maintains the genetic architecture of intercellular adhesion during chondrogenesis, a prerequisite for proper tissue morphogenesis.

### LAMA5 is required for pericellular matrix assembly and spheroid morphogenesis

Given that cell–cell junctions and ECM interactions together facilitate three-dimensional tissue organization, we next examined the morphological consequences of loss of LAMA5 using 3D chondrogenic spheroids. This model recapitulates essential features of cartilage development, as cells in spheroids undergo spontaneous condensation, establish junctional contacts, and secrete pericellular matrix (PCM) components that assemble into a stratified ECM. These processes collectively mimic the self-organizing behaviors and matrix deposition patterns observed during in vivo chondrogenesis, allowing quantitative assessment of tissue cohesion and morphogenesis.

During the chondrogenic spheroid differentiation process, we continuously monitored spheroids to observe for morphological differences between wildtype (WT) and KO spheroids. At day 5 of differentiation, brightfield imaging revealed striking morphological differences. WT cells formed compact, spherical aggregates, whereas KO spheroids appeared smaller, less cohesive, and irregularly contoured (Figure 3A). Quantitative morphometry confirmed a significant reduction in spheroid volume across all time points (days 5–20; Figure 3B), indicating persistent structural impairment. To dissect the dynamics of aggregate formation, we analyzed spheroid circularity over time. KO spheroids exhibited markedly reduced circularity during early stages (days 5–10), consistent with defective compaction and impaired mechanical symmetry (Figure 3C). Interestingly, this difference diminished by day 15 and reversed by day 20, suggesting time-dependent remodeling or compensatory structural changes.

**Figure 3:**
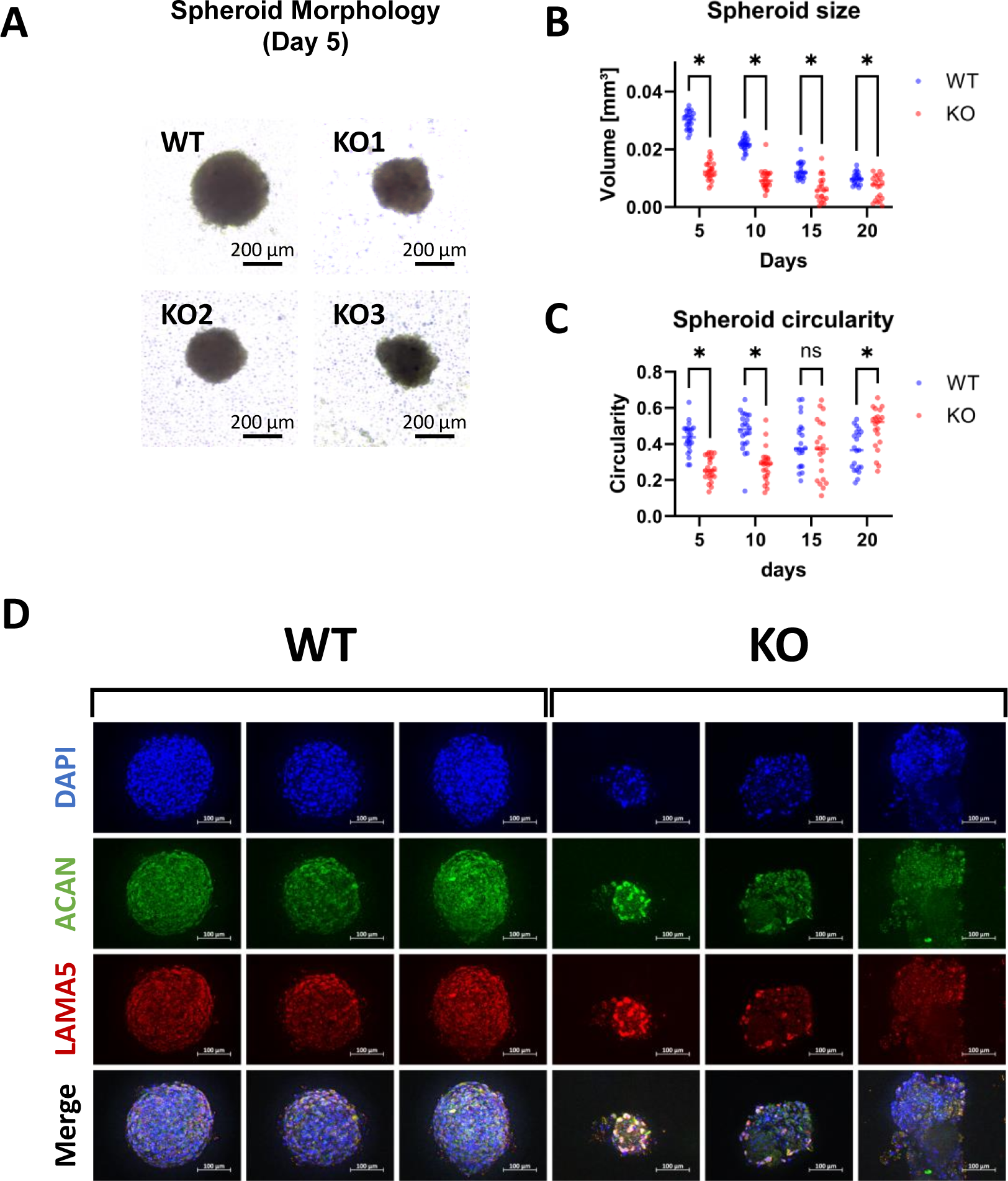
LAMA5 deficiency impairs chondrogenic spheroid morphogenesis. (A) Representative brightfield and images of WT and LAMA5-KO chondrogenic spheroids after 5 days of differentiation. WT spheroids display round aggregates, whereas KO spheroids remain smaller and irregular. (B) Quantitative morphometry confirmed a significant reduction in spheroid volume in KO cells at all timepoints (day 5 WT 0.0299 ± 0.003 mm³ vs KO 0.0127 ± 0.003 mm³, p < 0.0001; day 10 WT 0.0218 ± 0.002 mm³ vs KO 0.0096 ± 0.003 mm³, p < 0.0001; day 15 WT 0.0125 ± 0.003 mm³ vs KO 0.0063 ± 0.004 mm³, p < 0.0001; day 20 WT 0.0099 ± 0.002 mm³ vs KO 0.0068 ± 0.003 mm³, p = 0.0044; unpaired t-test with Welch correction, Holm–Šídák multiple-comparison adjustment, α = 0.05). (C) Circularity analysis revealed altered morphology at early stages (day 5 WT 0.434 ± 0.080 vs KO 0.262 ± 0.066, p < 1×10⁻L; day 10 WT 0.471 ± 0.106 vs KO 0.238 ± 0.091, p < 1×10⁻□), which normalized by day 15 (p = 0.39) but inverted by day 20 (WT 0.365 ± 0.112 vs KO 0.486 ± 0.115, p = 0.001). Data represent n = 21–24 spheroids per genotype and timepoint, pooled from three independent differentiations. (D) Immunofluorescence images of WT and LAMA5-KO chondrogenic spheroids after the 20-day differentiation period. WT spheroids display, round aggregates with continuous laminin α5 and aggrecan deposition, whereas KO spheroids display irregular shape and expression pattern. Scale bars: brightfield = 200 µm; immunofluorescence = 100 µm.

To reveal the cellular basis of these morphological phenotypes we performed immunofluorescence at day 20. WT spheroids displayed continuous pericellular laminin α5 and organized aggrecan deposition, hallmarks of an intact PCM (Figure 3D). In contrast, KO spheroids lacked pericellular laminin and exhibited fragmented, disorganized aggrecan architecture, correlating with their irregular morphology.

Together, these data demonstrate that LAMA5 is indispensable for PCM assembly and the physical integrity of chondrogenic spheroids. The impaired compaction and disrupted matrix organization in KO aggregates provide functional evidence that ECM–junction crosstalk is essential for cohesive tissue morphogenesis.

### LAMA5 loss rewires WNT-centered developmental programs rescued by LiCl

We next asked how the structural and junctional defects in LAMA5-KO cells translate into altered developmental signaling. Transcriptome-wide analyses across undifferentiated, osteogenic and chondrogenic differentiation states revealed coordinated downregulation of *WNT7A* and upregulation of *FLI1* in all KO cells (Figure 4A). These reciprocal changes were validated by RT–qPCR (Figure S1C, D) and were strongly inversely correlated across all samples (Figure 4B), suggesting a globally shared regulatory pathway impaired by LAMA5 loss.

**Figure 4:**
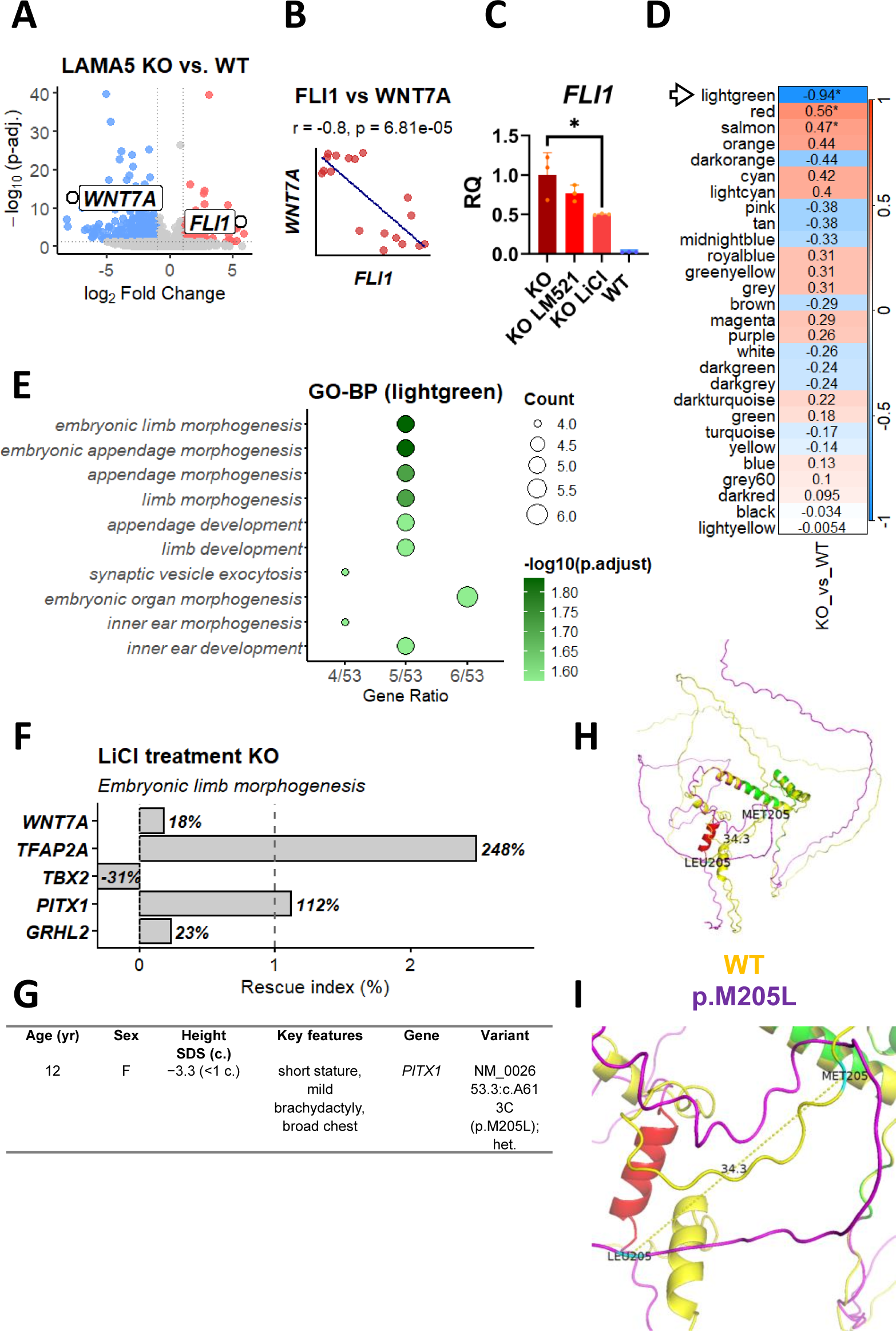
WGCNA identifies a skeletal development module with inversely regulated hub genes WNT7A and FLI1 and partial rescue by LiCl. (A) Volcano plot highlighting log2 fold-change of WNT7A and FLI1 across cell types (log2 fold-change WNT7A = −7.70, adjusted p = 2.18 x 10-16, n = 9; log2 fold-change FLI1 = +5.62, adjusted p = 3.48 x 10-7). (B) Correlation of FLI1 and WNT7A gene expression in all samples, revealing an inverse relationship (pearson r = -0.8, p = 6.8 x 10-5, Student’s t-test, total n = 18 samples). (C) Pharmacologic expression rescue; RT-qPCR of FLI1 in LAMA5-KO (Cell line 2) cells plated on LM521 (10 µg/ml, 7 days) or treated with LiCl (20 mM, 24 h). LiCl significantly shifted FLI1 expression toward WT levels (RQ KO = 1.00 ± 0.28; RQ LiCl = 0.50 ± 0.009, p = 0.031, Welch’s t-test, n = 3); LM521 produced a non-significant trend (RQ LM521 = 0.77 ± 0.10, p = n.s, Welch’s t-test, n = 3). (D) Module–trait correlation heatmap, the lightgreen module shows the strongest negative correlation with the KO trait (pearson r = −0.94, p < 0.05, Student’s t-test, total n = 18 samples). (E) GO Biological Process enrichment for the lightgreen module. Top terms include embryonic limb morphogenesis. (F) Rescue Index (%) highlighting partial restoration of gene expression in LAMA5-KO cells (cell line 2). Four (*WNT7A*, *TFAP2A*, *PITX1*, *GRHL2*) of the five genes from the KO-trait associated lightgreen module within the GO-term “embryonic limb morphogenesis” showed a trend towards expression restoration. (G) De-identified summary of a patient carrying a heterozygous PITX1 variant (NM_002653.3:c.A613C → p.M205L, VUS) who presented with isolated short stature. Variant confirmation by Sanger sequencing shown in Figure S3A; detailed clinical data in Table S1. (pctl. = percentile, VUS = Variant of Uncertain Significance). (H) Global structural overlay of WT (yellow) and mutant (magenta) PITX1 after rigid superposition (super in PyMOL). The canonical homeodomain (residues 89–148) is shown in green and the OAR motif (residues 280–293) in red. Note preservation of the homeodomain and misalignment of the OAR helix in the mutant. PyMOL v3.1.6.1 was used for visualization. (I) Close-up of residue 205 (cyan) showing the dashed Cα–Cα distance between WT and MUT residues (measured ≈ 34.3 Å) after superposition. Homeodomain (green) and OAR helix (red) are shown to provide spatial context.

Because canonical WNT signaling is central to cartilage and limb development, we tested whether pharmacologic activation of this pathway could rescue the observed defects. Treatment of KO cultures with lithium chloride (LiCl, 20 mM, 24 h), a GSK3 inhibitor that stabilizes β-catenin, partially normalized *FLI1* expression toward WT levels (Figure 4C). A similar, but weaker, trend was observed upon culture on recombinant laminin-521 (with the lacking α5-chain) for 7 days, indicating that both biochemical and structural interventions can modulate this pathway.

To understand whether the *WNT7A*–*FLI1* changes represented isolated events or part of a coordinated gene network, we performed weighted gene co-expression network analysis (WGCNA). This systems-level approach identifies modules of co-regulated genes based on similar expression patterns, allowing inference of shared biological pathways that might not emerge from differential expression alone. WGCNA revealed that *WNT7A* and *FLI1* clustered within the same gene expression module (light-green), which showed the strongest correlation with the KO genotype (Figure 4D, Figure S2A, B). We then performed GO-term analysis of this gene expression module, to investigate which pathways are affected by the genes most associated with the KO-trait. This revealed enrichment for the Gene Ontology term “embryonic limb morphogenesis,” indicating that LAMA5 loss perturbs a developmental gene network central to limb and cartilage formation (Figure 4E).

Based on the aforementioned gene expression rescue of *FLI1*, we next examined whether activation of canonical WNT signaling could also restore expression of genes within this KO-associated module. Using a “rescue index” derived from bulk RNA-seq (Figure 4F, Figure S2D), LiCl treatment shifted four of five embryonic limb-morphogenesis genes (*PITX1*, *GRHL2*, *TFAP2A*, and *WNT7A*) toward WT expression levels, whereas *TBX2* remained mostly unresponsive. At the network level, the top hub genes of the light-green module showed variable or reduced response to LiCl (Figure S2C, E), suggesting that short-term β-catenin stabilization could specifically benefit limb morphogenesis associated gene expression.

### Identification of PITX1 as a candidate for ISS

Notably, an internal clinical database search identified an unrelated individual with a heterozygous *PITX1* variant and isolated short stature (Figure 4G, Figure S3A, Table S1). *PITX1* is among the limb-morphogenesis genes that trended toward WT expression after LiCl treatment in our LAMA5-KO model, suggesting that perturbations at either extracellular (*LAMA5*) or transcriptional (*PITX1*) nodes may converge on the same developmental network. This observation provides preliminary clinical support that our observations are relevant to human growth phenotypes.

To assess whether the *PITX1* missense variant identified in a short-stature individual might alter PITX1 structure or function, we generated structural models of the wild-type and variant proteins and performed rigid and local superpositions (Figure 4H–I). The canonical PITX1 homeodomain (residues 89–148), which mediates sequence-specific DNA binding, remained structurally preserved in the mutant (green in Figure 4H), consistent with retention of the core DNA-binding fold. In contrast, the C-terminal OAR helix (residues 280–293; red in Figure 4H) was not aligned between WT and MUT models, indicating a local rearrangement of this regulatory motif. The mutated residue (Leu205) lies outside the homeodomain but in structural proximity to the OAR motif in our models and the Cα–Cα separation between WT and mutant residue 205 was measured as ∼34.3 Å after the reported superposition (Figure 4I). These data indicate that, although the DNA-binding domain is intact, the mutation is associated with altered conformation of a nearby regulatory module (OAR), which could modify PITX1 regulatory interactions with other proteins or intramolecular autoinhibition. Thus, even without disrupting DNA binding, the altered OAR configuration provides a plausible mechanism by which this variant could diminish PITX1 activity in pathways critical for skeletal and body growth.

These findings identify attenuation of canonical WNT signaling as a principal downstream consequence of LAMA5 deficiency and place *WNT7A* and *FLI1* within a disrupted developmental co-expression network. The partial, gene-specific rescue by LiCl implies that WNT pathway activation could counteract limb specific transcriptional consequences of ECM and junctional disruption. The subsequent identification of a short-stature patient with a heterozygous *PITX1* variant further supports convergence on this WNT-centered limb-morphogenesis module, linking our cellular findings to human growth phenotypes.

## Discussion

### LAMA5 deficiency disrupts ECM–junction programs while preserving lineage commitment

Previous studies have associated variants in ECM encoding genes, including *ACAN* and *COL2A1* with growth impairment^51,52^, but the contribution of laminins to lineage-specific differentiation and growth plate morphogenesis has remained largely unexplored. Our results position laminin α5 as a critical ECM regulator that links cell–cell adhesion, tissue morphogenesis, and developmental signaling in human chondrogenesis, providing mechanistic insight into ISS.

We therefore generated three independent LAMA5-KO USC clones, confirmed by Sanger sequencing, RT–qPCR, and western blot (Figure 1A), and validated that their differentiation potential remained intact (Figure 1B). Both WT and KO USCs robustly underwent osteogenic and chondrogenic differentiation, marked by differentiation associated gene expression (Figure 1C-F). This demonstrates that LAMA5 loss does not globally impair lineage commitment, which is consistent with reports that ECM perturbations can selectively affect morphogenetic organization without altering fundamental differentiation programs^53^. By maintaining baseline multipotency, this platform allows us to attribute observed morphogenetic and transcriptional defects specifically to laminin α5 deficiency, rather than generalized cellular dysfunction.

To get insights into the impact of LAMA5 impairment on the chondrogenic differentiation process, we performed bulk RNA-Seq of chondrogenic KO and wildtype cells (Figure 2). This revealed significant downregulation of *CDH1* (E-cadherin) and multiple claudin (CLDN) family members, indicating disruption of both adherens and tight junction assembly (Figure 1D). Mapping *CDH1* expression onto our previously established single-cell pseudotime atlas of USCs chondrogenesis^31^ demonstrated that its expression occurs during late-stage differentiation (Figure 1B), suggesting a failure to execute the terminal adhesion program. E-cadherin has been shown to cooperate with N-cadherin in human mesenchymal stem cell aggregates, promoting mesenchymal condensation and facilitating chondrogenic lineage commitment^54^. This highlights the critical role of *CDH1* in organizing condensation events, essential for proper tissue morphogenesis. These findings suggest laminin α5 as a key upstream ECM cue that sustains the transcriptional network governing intercellular adhesion, thereby providing a mechanistic link between ECM integrity, junctional organization, and the cellular cohesion necessary for growth plate formation and longitudinal skeletal growth^55^.

### 3D spheroid defects reveal early ECM–junction disruptions and biomechanical consequences

In order to provide further evidence to support this hypothesis, we analyzed three-dimensional (3D) chondrogenic spheroids (Figure 3). Here, knockout (KO) aggregates exhibited reduced size, irregular morphology, and diminished circularity during early differentiation stages (Figure 3A-C). Immunofluorescence analyses revealed fragmented aggrecan deposition and a lack of pericellular laminin α5, correlating with impaired spheroid compaction (Figure 3D). These phenotypic alterations provide functional evidence that laminin α5 stabilizes extracellular matrix (ECM)–junction interactions, which are essential for maintaining cohesive tissue architecture. Here, the observed time-dependent remodeling of KO spheroids suggests potential compensatory mechanisms (Figure 3C). However, these results underscore the critical role of early ECM–junction interactions in establishing proper cartilage morphology, a process that, when disrupted, is implicated in growth-plate dysfunction and growth deficits observed in idiopathic short stature (ISS)^29,56–58^. Furthermore, the observed fragmentation of aggrecan deposition and pericellular laminin in KO spheroids (Figure 3D) may compromise growth plate biomechanics. Aggrecan-rich PCM not only provides compressive resistance but also sequesters growth factors such as FGFs and BMPs, which are critical for growth plate chondrocyte proliferation and hypertrophy^8,59^. Loss of a continuous laminin–aggrecan network could therefore reduce local signaling efficiency, compounding transcriptional disruptions and resulting in structural immaturity. Importantly, the early compaction and PCM defects we observe in LAMA5-KO spheroids provide a plausible mechanistic basis for altered developmental signaling. Pericellular ECM organization and junctional integrity are known to modulate WNT/β-catenin responsiveness and β-catenin localization, such that changes in ECM composition or mechanical cues can alter WNT output^60^.

### Laminin α5 regulates canonical WNT signaling and embryonic limb-morphogenesis networks

Consistent with this model, transcriptomic profiling revealed that loss of LAMA5 rewires WNT signaling, as evidenced by downregulation of *WNT7A* (Figure 4A). Canonical WNT/β-catenin signaling is well-established as a critical regulator of chondrocyte proliferation, hypertrophy, and growth plate organization^61–63^. The coordinated dysregulation of *WNT7A* and *FLI1* in LAMA5-KO cells, along with their integration within a WGCNA-derived module enriched for embryonic limb morphogenesis genes (Figure 4D, E), suggests that laminin α5 contributes to the spatial and temporal organization of these signaling cascades. In further experiments, pharmacologic activation of WNT signaling via LiCl not only restored expression of *FLI1*, but also of (4/5) embryonic limb development-associated genes *PITX1*, *GRHL2*, *TFAP2A*, and *WNT7A* (Figure 4F), indicating that the transcriptional consequences of ECM disruption can be mitigated by downstream pathway modulation. These observations align with prior studies showing that ECM composition can influence WNT responsiveness and β-catenin localization, suggesting a functional ECM–signaling crosstalk^64–66^. Notably, LiCl treatment activated the embryonic limb-morphogenesis module across the board, including *TBX2*, which in our dataset was already elevated in LAMA5-KO cells and was further induced by LiCl (Figure S2D). Given that LiCl acts via GSK3 inhibition to stabilize β-catenin^67,68^, and that TBX2/TBX3 act downstream of canonical WNT signaling^69^, the concurrent induction of TBX2 is consistent with pathway activation rather than failed rescue. It is therefore plausible that LiCl restores the limb-morphogenesis network globally, and that the TBX2 increase represents pathway activation superimposed on an earlier compensatory upregulation in KO cells.

### Convergence with a clinical PITX1 variant supports a shared developmental axis in ISS

The clinical significance of our research is highlighted by the identification of a patient with isolated short stature who harbours a heterozygous *PITX1* missense variant (Figure 4G). *PITX1*, a key limb-patterning transcription factor^70^, resides in the limb-morphogenesis module perturbed by LAMA5 deficiency and showed partial restoration upon LiCl-mediated WNT activation. This raises the hypothesis that extracellular matrix disruption (*LAMA5*) and transcriptional dysregulation (*PITX1*) can perturb a common WNT-centered developmental axis, offering a unifying framework for otherwise heterogeneous short-stature phenotypes. Although preliminary, this single case provides a clinical anchor for the mechanistic pathway uncovered in our model.

Unlike previously reported *PITX1* mutations that predominantly cause complex limb malformations including clubfoot, preaxial polydactyly, and tibial deficiencies (e.g., E130K in the homeodomain^71^ or frameshifts disrupting the OAR domain^72^), the L205 variant identified here is associated with an isolated idiopathic short stature phenotype without limb malformations. Our structural models show preservation of the canonical homeodomain and DNA-binding fold, with structural domain alterations localized to the OAR motif and the adjacent residue 205 (Figure 4 H, I). This may lead to subtler alterations in PITX1’s transcriptional regulation capacity rather than gross developmental defects. Such a hypomorphic effect could selectively impair growth-plate function or the regulation of genes required for osteogenic and chondrogenic differentiation, providing a plausible molecular basis for the isolated short-stature phenotype. Supporting this, Pitx1 knockout mice show craniofacial and skeletal hypoplasia, including missing hyoid bone and delayed fontanelle closure, along with short stature, consistent with a role for PITX1 in growth regulation beyond limb identity^73^. Thus, the molecular location and nature of the *PITX1* variant may determine the phenotypic spectrum, with L205 producing a milder, isolated growth phenotype.

### Implications for ECM-driven growth disorders and therapeutic targeting

In summary, our work establishes a mechanistic link between extracellular matrix composition, junctional organization, and canonical WNT signaling in human chondrogenesis, positioning laminin α5 as a critical regulator of limb morphogenesis and growth plate function. The partial rescue of gene expression defects by pharmacologic WNT activation using LiCl highlights the therapeutic potential of targeting this pathway in ECM-related growth disorders. Moreover, the identification of a novel *PITX1* variant associated with isolated short stature, distinct from classical limb malformation phenotypes, expands the genetic landscape of idiopathic short stature and underscores the convergence of extracellular and transcriptional defects on a shared developmental network. These findings provide novel insights for the diagnosis and intervention in growth disorders and warrant further investigation into *PITX1* as a candidate gene in ISS.

## Data and code availability

The datasets and code supporting this study are in the process of being deposited in a public repository but are not yet publicly available due to ongoing curation and formatting. They are available from the corresponding author upon reasonable request.

## Acknowledgments

This work was funded by the Deutsche Forschungsgemeinschaft (DFG, German Research Foundation) – [TH896/7-1]. We thank all the patients and their families for participating in this project. We want to thank the FAU FACS core unit NFZ for cell sorting and advice. Additionally, we want to thank Evelyn Galsterer for excellent technical assistance.

## Author contributions

A.S: performed the experiments, processed and analyzed the data, and wrote the original draft, E.B: processed the raw sequencing data. S.U: uploaded all sequencing data, A.B.E: assisted with sample collection, C.T.T: project P.I, acquired funding, supervised research and study design, and revised manuscript

## Web Resources

AlphaFold Protein Structure Database, https://alphafold.ebi.ac.uk/

ColabFold, https://colabfold.mmseqs.com

OMIM, http://www.omim.org

UCSC BLAT, https://genome.ucsc.edu/cgi-bin/hgBlat

