## Supplemental Information for "LAMA5 deficiency disrupts ECM–WNT crosstalk in chondrogenesis and contributes to idiopathic short stature"

### Supplemental Figures

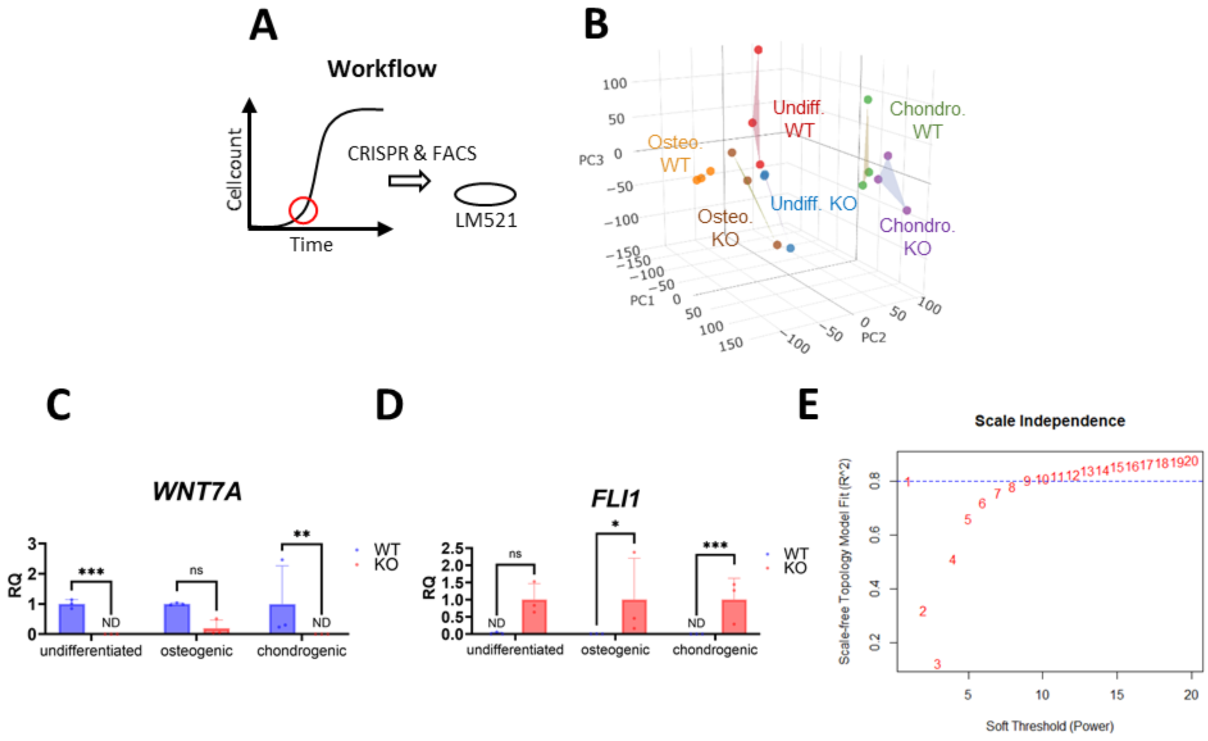


**Figure S1.** A: Cartoon (cell counts / time) for USCs illustrating characteristic S-shape kinetics and the transient rescue of LM521 coating during expansion. B: Principal component analysis (PCA), revealing clear separation regarding genotype and cell type underscoring general uniqueness of samples: C/D: Confirmation of differential gene expression of the genes *WNT7A*/*FLI1* with RT-qPCR in all cell types. C: WNT7A expression (Undifferentiated: RQ WT = 1.00 ± 0.15, RQ KO = $8.8 \times 10^{-5} \pm 4.7 \times 10^{-5}$, p = $7.6 \times 10^{-4}$, Welch's t-test, n = 3; Osteogenic: RQ WT = 1.00 ± 0.04, RQ KO = 0.20 ± 0.27, p = n.s., Welch's t-test, n = 3; Chondrogenic: RQ WT = 1.00 ± 1.27, RQ KO = $7.3 \times 10^{-4} \pm 1.1 \times 10^{-3}$, p = $8.0 \times 10^{-3}$, Welch's t-test, n = 3). D: FLI1 expression (Undifferentiated: RQ WT = 0.021 ± 0.035, RQ KO = 1.00 ± 0.47, p = n.s., Welch's t-test, n = 3; Osteogenic: RQ WT = 0.0068 ± 0.0007, RQ KO = 1.00 ± 1.20, p = 0.029, Welch's t-test, n = 3; Chondrogenic: RQ WT = $4.0 \times 10^{-4} \pm 2.0 \times 10^{-4}$, RQ KO = 1.00 ± 0.62, p = $8.8 \times 10^{-4}$, Welch's t-test, n = 3). E: WGCNA scale-free topology plot (pickSoftThreshold), showing scale independence ($R^2$) vs soft threshold power (1–20) with the chosen power = 9.

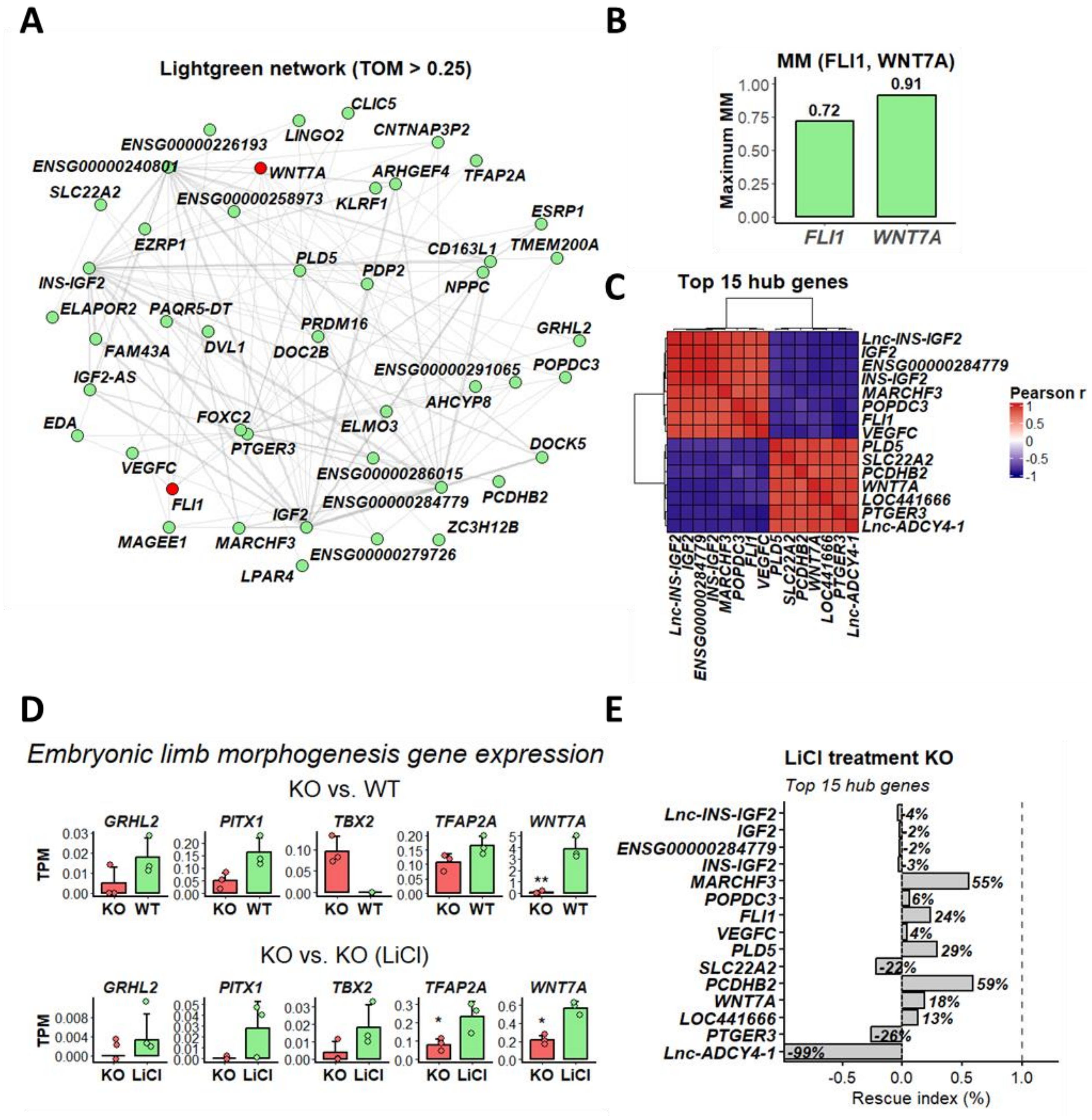


**Figure S2.** A: WGCNA network of the light green module, filtered for topological overlap measure (TOM) > 0.25 (FLI1 and WNT7A highlighted in red). B: Module membership (kME) for WNT7A and FLI1 with strong membership in the "lightgreen" module (kME WNT7A = 0.91; kME FLI1 = 0.72). C: Pairwise correlation matrix for the top 15 hub genes of the lightgreen module. F: Barplots depicting gene expression (TPM) of the five genes from the KO-trait associated lightgreen module within the GO-term "embryonic limb morphogenesis". Here, the two bulk RNA-Sequencing batches KO (all cell lines) vs. WT and KO (cell line 2) vs KO (cell line 2 + 10 mM LiCl) were not merged. Annotated significance stars represent adjusted p values from DESeq2 within the batch (adjusted $p < 0.05$ = *, adjusted $p < 0.01$ = **). E: Rescue Index (%) highlighting changes in top 15 hub gene expression upon LiCl treatment.

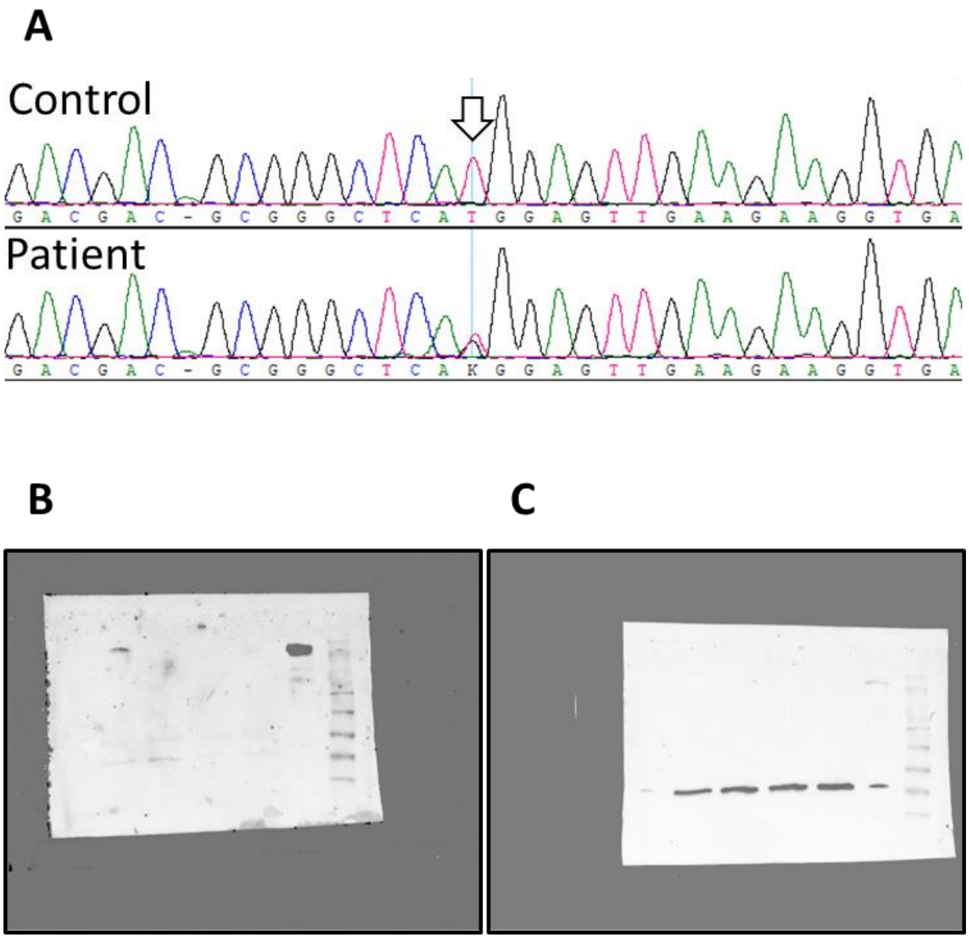


**Figure S3.** A: Electropherogram confirming the heterozygous *PITX1* variant identified by exome sequencing (reverse complementary). The variant base is indicated by arrows. B/C: Unedited westernblot images from LAMA5 (A) and Actin beta (B), as acquired from the ChemiDoc imaging system (bio-rad)

**Table S1:** Clinical and genetic summary of a de-identified patient with isolated short stature carrying a heterozygous PITX1 variant (NM_002653.3:c.A613C → p.M205L). Variant confirmed by Sanger sequencing (Supplementary Figure 3A).

| Field | Entry |
|---|---|
| **Sex** | F |
| **Age at examination** | 12 years |
| **Height (cm)** | 131.4 cm (−3.3 SDS; <1st centile) |
| **Weight / BMI** | 36.2 kg / 18.3 kg/m² |
| **Head circumference** | 51 cm (29th centile) |
| **Key phenotypic features** | Mild brachydactyly; broad chest; mildly low-set ears; short/broad forefeet; otherwise normal psychomotor development |
| **Birth history** | 38 wks; BW 2685 g; APGAR 9/10/10 |
| **Neurological / audiological findings** | Sensorineural hearing impairment (familial) – not subject of current study |
| **Karyotype / CNV** | 46,XX; no pathogenic CNV detected |
| **Test method** | Trio exome sequencing (panel of 2,175 short-stature genes) + Sanger confirmation |
| **Genomic coordinates (hg38)** | chr5:135029111 T > G |
| **Gene / Transcript** | *PITX1* (NM_002653.3) |
| **Variant (HGVS)** | c.A613C (p.M205L) |
| **Exon / Type** | Exon 3; missense (nonsynonymous SNV) |
| **Zygosity** | Heterozygous |
| **Confirmation** | Sanger sequencing (see Supplementary Figure 3A) |
